# Investigating the role of cortical microglia in a mouse model of viral infection-induced seizures

**DOI:** 10.1101/2025.08.22.671562

**Authors:** Lakshmini Balachandar, Lauren Buxton, Ireland Kearns, Matthew F. Stefanic, Laura A. Bell, Ana Beatriz DePaula-Silva, Karen S. Wilcox

**Affiliations:** Department of Pharmacology and Toxicology, University of Utah, Salt Lake City, US

**Keywords:** Microglia, Epilepsy, Cortex, Theiler murine encephalomyelitis virus (TMEV), Viral infection, Encephalitis, Calcium imaging, Purinergic signaling, Cytokines

## Abstract

Microglia, resident immune sentinels in the brain, are crucial in responding to tissue damage, infection, damage signals like purines (ATP/ ADP), and clearing cellular debris. It is currently unknown how microglial reactivity progresses and contributes to seizure development following Theiler’s Murine Encephalomyelitis Virus (TMEV) infection. Previously, our group has demonstrated that purinergic signaling in microglia is disrupted in the hippocampus of TMEV-infected mice. However, whether reactive cortical microglia also exhibit changes in purinergic signaling, cytokine levels, and purinergic receptors are unknown. Thus, we seek to evaluate region-based differences in microglial reactivity in the TMEV model. We employed a custom triple transgenic mouse line expressing tdTomato and GCaMP6f under a CX3CR1 Cre promoter and exogenously applied ATP/ADP to acute brain slice preparations from TMEV-infected mice and controls. Interestingly and in contrast to what is observed in hippocampus, we found that despite microglial reactivity in the cortex, microglia can respond to purinergic damage signals and engage calcium signaling pathways, comparable to PBS controls. Using a cytokine panel, we also found that pro-inflammatory cytokine levels (TNF-α, IL-1α and IFN-γ) are brain-region dependent in mice infected with TMEV. Using RNAScope-FISH, we observed increases in expression of purinergic receptors responsible for microglial motility (P2Y_12_R) and inflammation (P2X_7_R) in the cortex. Collectively our results suggest that following TMEV infection, microglial response to novel damage signals, as well as the production of proinflammatory cytokines, varies as a function of brain region.

## INTRODUCTION

Temporal lobe epilepsy (TLE), a severe and devastating seizure disorder, is one of the most prevalent forms of acquired epilepsy and the most challenging to treat with current antiseizure drugs (Engel Jr, 2001). Central nervous system infections, including those caused by neurotropic viruses, are a significant cause of TLE. Several known viruses including Human Herpes simplex virus (HHV), varicella-zoster virus (VZV), enteroviruses, Japanese Encephalitis Virus, West Nile Virus and Human Immunodeficiency Virus cause acute clinical seizures and subsequent development of TLE (Misra et al., 2008; Singhi, 2011). Despite the prevalence of viral infections leading to epilepsy, there currently is no therapeutic approach for successful prevention of epileptogenesis in patients. The Theiler’s Murine Encephalomyelitis Virus (TMEV) mouse model of infection-induced temporal lobe epilepsy (TLE) is an invaluable tool in epilepsy research for investigating novel therapeutic approaches to prevent viral-infection induced seizures in patients (DePaula-Silva et al., 2021). Mice (C57Bl6/J) infected with the Daniel’s strain of TMEV display both spontaneous and handling-induced seizures during the acute phase following infection (3-8 dpi) (Libbey et al., 2008; Patel et al., 2017). The mice develop significant neuronal loss, predominantly in the CA1 and CA2 regions of the hippocampus, persistent gliosis across several brain regions i.,e., hippocampus, cortex, and limbic system (Bell et al., 2020; Loewen et al., 2016; K.-A. A. Stewart et al., 2010) and dramatic increases in expression of proconvulsant cytokines and reactive oxygen species (Bhuyan et al., 2015; Kirkman et al., 2010; Patel et al., 2017; Umpierre et al., 2014). Several weeks later, seizure thresholds are reduced and the majority of animals that demonstrated acute seizures develop TLE and behavioral comorbidities (Bröer et al., 2016; K.-A. A. Stewart et al., 2010; K. A. A. Stewart et al., 2010; Umpierre et al., 2014). Seizures likely originate in the hippocampus and over the course of infection, begin to secondarily generalize to the cortex (Patel et al., 2017).

Microglia are resident immune cells in the central nervous system (CNS) (Schafer et al., 2013) that play a crucial role in responding to tissue damage and infection. Microglial reactivity constitutes their innate immune response to pathogens, damage signals and cellular debris, but chronic persistence of this reactivity can lead to exacerbation of the neuroinflammatory milieu (Eyo et al., 2017; Henning et al., 2023; Illes, 2020; Illes et al., 2020). During homeostatic condition, continuous microglial surveillance in the brain (Nimmerjahn et al., 2005) is accompanied by spontaneous calcium transients (Brawek & Garaschuk, 2013). The release of damage signals in the brain environment induces calcium transients in microglia, orchestrating a host of downstream second messenger cascades and ultimately leading to their reactivity and response to novel damage cues and aberrations in neuronal activity (Damisah et al., 2020; Hughes & Appel, 2020). Following inflammatory insults, they have been reported to have robust increases in calcium transients, accompanied by directional process movement (Brawek et al., 2014; Eichhoff et al., 2011; Pozner et al., 2015; Umpierre et al., 2020). Following CNS infection and injury, reactive microglia along with infiltrating macrophages, contribute to seizure activity (Cusick et al., 2013; Wilcox & Vezzani, 2014). In TMEV-infected mice, these cell types initiate a cascade of the innate immune responses in the CNS, initiating the generation of a cytokine storm encompassing high levels of production of pro-convulsant cytokines like TNF-α, IL-1β, IL-6 and IFN-γ and chemokines. This leads to amplified calcium dynamics during infection and has an effect on seizure activity (Henning et al., 2023; Hide et al., 2000; Wilcox & Vezzani, 2014). Additionally, reactive microglia also serve as a major source of TNFα in mice with TLE (Henning et al., 2023).

Seizures also induce microglial interactions with neuronal somata and dendrites, including via purinergic receptors. Microglial migration and chemotaxis are heavily dependent on purinergic receptors including the P2Y_12_ receptor, which is predominantly expressed in microglia (Dissing-Olesen et al., 2014; Eyo et al., 2014; Gómez Morillas et al., 2021). Specifically, reactive microglia in the TMEV model have been reported to have significant gene expression changes including downregulation of P2Y_12_R (DePaula-Silva et al., 2019; Wallis et al., 2024). Reactive microglia also exhibit morphological and functional changes, differential expression of surveillance genes governing damage signal recognition, and cytoskeletal reorganization and purinergic receptors including P2X_7_R and P2Y_12_R (DePaula-Silva et al., 2019; Hammond et al., 2019; Jimenez-Pacheco et al., 2013; Lively & Schlichter, 2018). Functional upregulation of P2X_7_R has been observed in microglia *in vivo* in human and rodent TLE models and following lipopolysaccharide (LPS) injections, high levels of ATP, and upregulation of pro-inflammatory cytokines (Choi et al., 2007; He et al., 2017; Jimenez-Pacheco et al., 2013). P2X_7_R is a key regulator of exacerbating the inflammation cascade via the nucleotide-binding domain, leucine-rich–containing family, pyrin domain–containing-3 (NLRP3) inflammasome, and thereby acting as a mediator in initiating caspase cascades involved in apoptosis (Di Virgilio et al., 2017; Suzuki et al., 2020). It has also been shown that P2X_7_R antagonism led to reduction in spontaneous seizures and gliosis in TLE (Jimenez-Pacheco et al., 2016).

Despite recent advances, there is a major gap in understanding the mechanistic and region-specific role of reactive microglia in seizure development in this infection induced model. In a recent study from our lab, during the peak period of acute TMEV infection, reactive hippocampal microglia had disrupted calcium signaling responses to local purinergic insults/ puffing and exhibited dampened motility towards laser-burn damage in acute brain slice preparations. Interestingly, P2Y_12_R was downregulated in the hippocampus, which could have led to observed motility deficits and microglial TNF-α upregulation in the hippocampus of TMEV-infected mice (Wallis et al., 2024). Since the cortex is involved in generalized seizures in the TMEV model (Patel et al., 2017), whether reactive cortical microglial calcium signaling and responses to damage cues could lead to production of pro-inflammatory cytokines, changes in cytoskeletal receptors and eventually contribute to seizure propagation, is yet to be investigated.

In this study, we employed a multidisciplinary approach to assess, during the peak TMEV-infection period, (i) the ability of cortical microglia to respond to purinergic damage signals via calcium signaling in acute brain slice preparations, (ii) brain region-specific production of pro-inflammatory cytokines/ chemokines, and (iii) changes in purinergic receptor expression in reactive microglia. Interestingly, we found that reactive cortical microglia can respond to novel purinergic damage cues (ATP and ADP) comparable to PBS controls, during the peak period of TMEV infection (5 dpi). We also evaluated brain-region dependent cytokine and chemokine levels in the TMEV model to better understand regional responses to infection, specifically in the cortex versus hippocampus. We observed an increase in levels of inflammatory markers like TNF-α, IL-1α, IFN-γ and IFN-β, amongst others, in the cortices of TMEV mice, as compared to PBS controls, which were further increased in the hippocampus. Using RNAScope-FISH, we also observed a significantly higher expression of P2Y_12_R, P2X_7_R and TNF-α in cortical microglia post TMEV-infection. Overall, cortical microglia, during the acute peak period of TMEV infection, retain their ability to respond to novel purinergic damage cues, despite being reactive, and having enhanced purinergic receptor and cytokine expression as compared to PBS controls. Following TMEV infection, there is reduced expression of cytokine and chemokine profiles in the cortex as compared to the hippocampus. These findings pave the way for future investigation on the impact of regional gene expression changes in reactive microglia in seizure generation.

## MATERIALS AND METHODS

The experimental procedures performed as part of this study were carried out in compliance with the National Institutes of Health Guide for the Care and Use of Laboratory Animals and approved by the Institutional Animal Care and Use Committee (IACUC) at the University of Utah. The timeline of experiments and various techniques employed in this study are shown in Figure 1.

**Figure 1.**
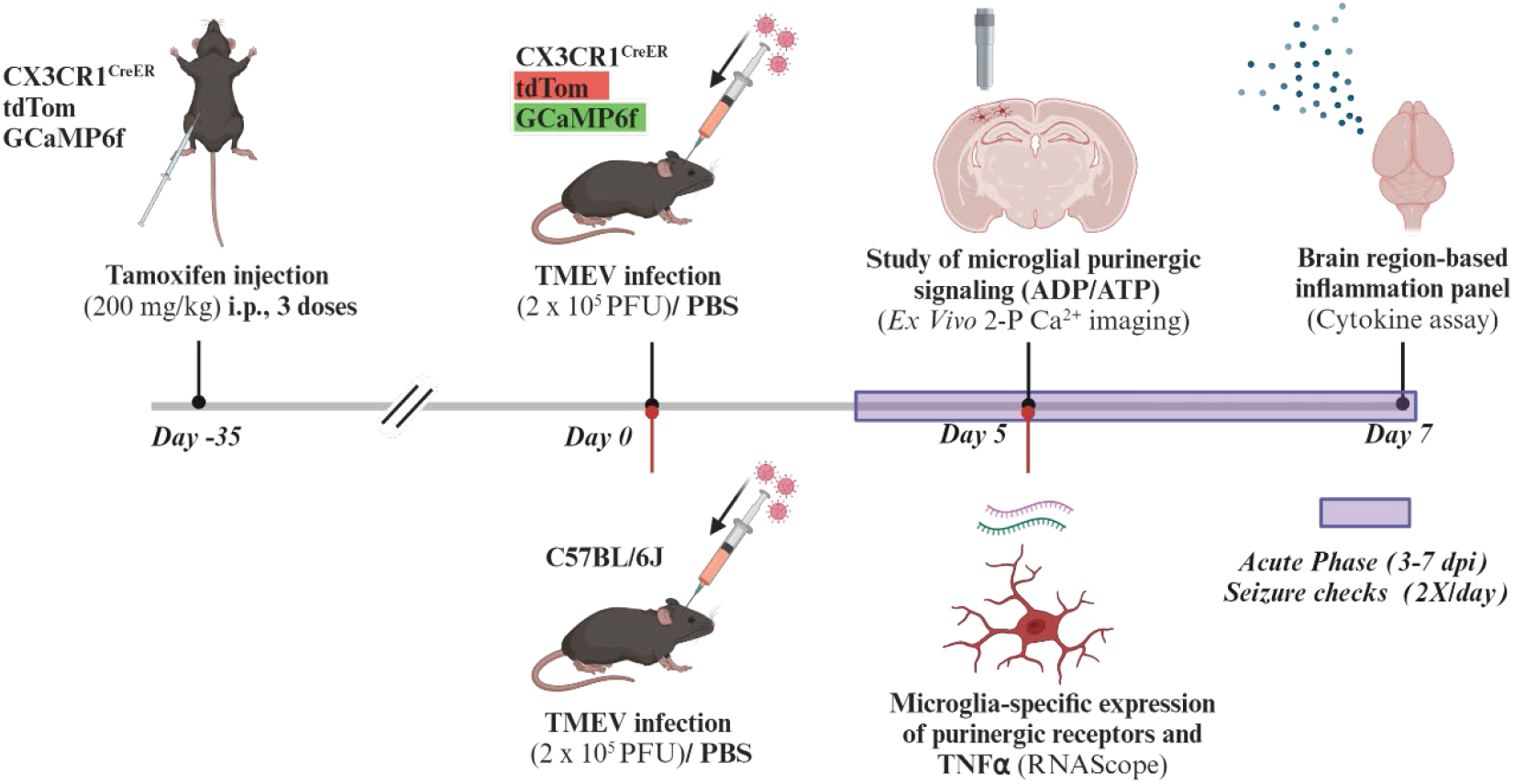
Timeline of tamoxifen administration, TMEV infection in mice, seizure monitoring, cellular, molecular, and calcium imaging experiments performed in this study. Mice heterozygous for CX3CR1-tdTomato-GCaMP6f were injected with tamoxifen (i.p.) to induce expression of tdTom and GCaMP6f. After at least 35 days to allow for macrophages to turn over and not express the transgenes, mice were injected intracranially with TMEV or PBS and were monitored twice a day from 3-7 days for seizures. Acute brain slices were made from mice at 5 dpi for 2-P calcium imaging. At 7 dpi in a separate cohort, mice brains were collected for the cytokine assay. Bottom panel of timeline: C57BL/6J (naïve) mice were injected with TMEV or PBS intracranially and were monitored twice a day from 3-5 days for seizures. At 5 dpi, mice brains were prepared for RNAScope analysis. Figure created using Biorender.

### Animals

Mice were housed in standard cages, provided with food (Teklad Global Soy Protein-Free Extruded Rodent Diet cat. #2920X; Harlan Laboratories) and water *ad libitum* in a 12h-12h light-dark cycle. All experiments were randomized and performed in both female and male mice. Custom triple transgenic mice heterozygous for CX3CR1, tdTomato (tdTom) and Lck-GCaMP6f were generated from Jackson (JAX) lab mice lines (020940, 007914 and 029626) at our vivarium. Expression of all transgenes were confirmed by PCR. These transgenic mice were used for experiments involving acute brain slices and cytokine assays. For RNAScope experiments, wild type C57BL/6J mice (JAX labs 000664) were used. After arrival, the mice were acclimatized to our animal facility and diet for at least 1 week before experiments.

### Tamoxifen-induced recombination

Transgenic mice (older than 5 weeks of age) were administered with three doses of tamoxifen (TAM) (Sigma-Aldrich T5648) dissolved in peanut oil (20 mg/ml) (i.p., 200mg/kg, every 48 hours) to induce Cre-mediated gene expression of tdTomato and Lck-GCaMP6f. The use of CX3CR1 creERT2/+ mice meant that infiltrating macrophages would also express GCaMP6f and tdTom thereby making it difficult to distinguish from resident microglia (Cusick et al., 2013). Therefore, subsequent treatments were conducted at least thirty-five days after the last tamoxifen injection, so that infiltrating macrophages would not display the tamoxifen-induced fluorophores (Parkhurst et al., 2013).

### TMEV infection and seizure monitoring

Mice were injected with 20uL of either 2.5 x 10^5^ PFU of Daniel’s strain of TMEV or phosphate buffered saline (PBS) intra-cortically, 2mm deep in the right hemisphere of the posterior parietal cortex, under isoflurane anesthesia and compressed air (Loewen et al., 2016; Patel et al., 2017; K.-A. A. Stewart et al., 2010). Mice were checked for handling-induced behavioral seizures twice a day, starting at 3 dpi as previously described. The seizures were scored based on a modified racine scale: stage 3, forelimb clonus; stage 4, additional rearing; stage 5, additional rearing and falling, few jumps; and stage 6, additional clonic running, extensive jumping, falling and severe hindlimb clonus (Patel et al., 2017; Racine, 1972). Only mice that have had at least one observed seizure above stage 3 were used for experiments.

### Acute brain slice preparation

All *in vitro* experiments were performed during the peak period of TMEV infection (5 dpi) in the cortex, on the ipsilateral side of infection, in mice aged 10-20 weeks. At 5 dpi, mice were anesthetized using isoflurane anesthesia and oxygen. Once the mouse lost its righting reflex, it was rapidly decapitated, the brain was extracted and placed in ice-cold cutting N-methyl-D-glucamine (NMDG) solution for 10 seconds. The hind brain was trimmed and was mounted on the vibratome (Vibratome 3000, Vibratome Company) with a back-support of a 4% agarose (wt/vol) block while making sections. Coronal sections (350μm) containing the cortex were made using the NMDG cutting solution (92 mM NMDG, 2.5 mM KCl, 30mM NaHCO_3_, 1.2 mM NaH_2_PO_4_, 20 mM HEPES, 25 mM Glucose, 5 mM Sodium Ascorbate, 2mM Thiourea and 3mM Sodium Pyruvate, 0.5 mM CaCl_2_,10mM MgSO_4_). Sections were incubated at 32-33°C for 30 minutes with regular 2000 mM NaCl spike-ins depending on age of the mouse (Ting et al., 2018). Subsequently, slices were transferred to room temperature (RT) HEPES holding solution (92 mM NaCl, 2.5 mM KCl, 30 mM NaHCO_3_, 1.2 mM NaH_2_PO_4_, 20 mM HEPES, 25 mM Glucose, 5 mM Sodium Ascorbate, 2mM Thiourea and 3mM Sodium Pyruvate, 2 mM CaCl_2_,10mM MgSO_4_) and held until imaging. All solutions were constantly bubbled with carbogen (95% O_2_/5% CO_2_), pH titrated to 7.35 +/-0.05, and osmolarity at 290-310 mOsm). The reagents used for solution preparation were purchased from Sigma-Aldrich. The slices were imaged for up to 6 hours after sectioning.

### Two-photon calcium imaging in microglia

Brain slices were placed in a custom slice chamber and hold-down (W4 64-0249, Warner Instruments) to prevent movement during imaging. Slices were continuously perfused with artificial cerebrospinal fluid (aCSF) (126 mM NaCl, 3 mM KCl, 26 mM NaHCO_3_, 1.4 mM NaH_2_PO_4_, 10 mM Glucose, 2 mM CaCl_2_,12 mM MgSO_4_) by a peristaltic pump and bubbled with carbogen (95% O_2_/5% CO_2_). The temperature of the bath was maintained at 24-26°C using an inline heater (TC-324C, Warner Instruments). Two-photon (2-P) calcium imaging was performed on a Prairie Ultima system (Bruker Corporation) using a Mai Tai DeepSee EHP 1040 laser (Spectra Physics) at 69 mW laser power, Prairie View software, a 20X water-immersion lens (NA: 1, Olympus), and emission bandpass filter at 560 nm to split green from red wavelengths (Bruker 370A510816). GCaMP6f and tdTom constitute the green and red channels respectively, and images of microglia were acquired at 920 nm excitation, where both fluorophores were excited optimally. Microglial calcium imaging was performed ∼50 µm from the surface of the slice to exclude tissue damage due to slicing. ATP and ADP working solutions were prepared from stocks (10 mM ATP (Tocris 3245)/ ADP (Tocris 1624) in reverse osmosis (RO) water, stored at −80 °C). The stocks were diluted to 100 µM ATP or ADP in aCSF with 15 µg/mL Alexa568 (Invitrogen A33081) to visualize the puff. Puff pipettes were pulled by a HEKA PIP 6 electrode puller from 1.5 mm OD, thin-walled borosilicate glass and had an open tip resistance of 2-3.5 MΩ. ATP was dispensed using a Picospritzer III system (Parker Instrumentation) with 6 PSI pressure for 350 ms. The ipsilateral side receiving the TMEV or PBS injection was imaged in the cortical regions of layers II-V and either ATP or ADP was applied to different fields of view in the same brain slice. Time series images were acquired at 920 nm excitation, 2 Hz, 1.2 µs/pixel dwell, 512 x 512 pixels per frame, 2.5x optical zoom, and 240 pockels laser power for a 13s baseline and 1 min after the puff.

### Detection of changes in calcium responses in response to ATP/ ADP application

To quantify changes in microglial calcium response to ATP and ADP puffs, image noise was reduced with a hybrid 3D median filter in ImageJ (Schindelin et al., 2015). The area of ATP agonist spread in the brain slice was identified by the spread of Alexa 568 in the post-application period, as previously described in our lab (Umpierre et al., 2019). Next, the area of the puff was demarcated, and a mask was created in order to facilitate detection of cells within the puff. Using CellProfiler (Stirling et al., 2021) cell image analysis software, various microglial cells were identified and segmented, rendering them as regions of interest (ROIs). This output was further analyzed on ImageJ, and measures of the ROIs were calculated from the mean pixel intensity in the ROI (F) for each point in time. Using a custom MATLAB (MathWorks, Natick, MA, USA) script, the maximum of calcium signal change (F-F_0_)/F_0_ was calculated compared to the mean pixel intensity for baseline 13s before the application (F_0_). The baseline was defined as the mean of the fluorescent intensity values before the puff. The event threshold is set to two standard deviations above the initial baseline fluorescence and the “findpeaks” feature was used. ΔF/F_0_ time-series plots were generated by averaging F values of each pixel at each time point and using the mean fluorescence of all image frames as the baseline fluorescence (F_0_).

### Cytokine and chemokine panel

TMEV-infected and PBS-control mice (aged 12-16 weeks) were euthanized at 7 dpi by transcardial perfusion with PBS, and the ipsilateral hippocampi and cortices were dissected, and flash frozen in liquid nitrogen. The tissue was homogenized using a mechanical homogenizer in ∼50 μL 1X PBS. Furthermore, the tissue was lysed with a lysis buffer (R&D Systems 895347) and protease inhibitor cocktail (Roche, 04693159001), centrifuged, following which the lysate was extracted for protein quantification using the BCA assay kit (ThermoFisher Scientific 23225). Normalized protein amounts across samples were loaded to perform the cytokine assay. Cytokine and chemokine levels were measured using the LEGENDplex™ Mouse Inflammation Panel (Biolegend 740446), according to manufacturer’s protocol. Briefly, standards were prepared by serially diluting standards provided in the kit (used for calibration of analyte curves). Assay buffer (25 μL) and samples / standards (25 μL) were added to wells of a 96 well V-bottom plate. 25 μL of mixed beads were added after vortexing them (to avoid bead settling). Samples were shaken at 80 rpm on a plate shaker for 2 hours at RT, centrifuged at 1050 rpm for 5 minutes and the supernatant was discarded in one continuous and forceful motion. The centrifugation step was repeated once more, and 25 μL biotinylated detection antibodies were added to each well, followed by shaking at 800 rpm for 1 hour at RT. 25 μL of streptavidin-PE (SA-PE) was added to each sample and incubated for 30 minutes. Finally, samples were washed with the wash buffer and the beads were resuspended. Samples were subjected to flow cytometry analysis (BD CytoFlex), on the same day as the assay. Data analysis was performed using BioLegend’s LEGENDplex™ data analysis software and statistical analysis was carried out on GraphPad Prism (version 9.4).

### RNAScope in situ-hybridization and immunohistochemistry

TMEV-infected and PBS-control mice (aged 10-12 weeks) were euthanized at 5 dpi using excessive isoflurane and transcardially perfused with 1X PBS briefly for ∼30 seconds (until liver was ∼75% clear to preserve RNA quality), followed by 10% neutral buffered formalin solution (NBF). The brains were postfixed for 24 hours in 10% NBF and subsequently transferred to a 15%/30% sucrose gradient for cryoprotection. Coronal sections of the brain were made (15 µm thickness) using a freezing stage microtome (Leica SM 2010R, Buffalo Grove, IL). Duplicate sections from each brain were mounted on Superfrost slides (Fisher Scientific, 1255015) and processed for RNAScope (controls included). Fluorescent in situ hybridization (FISH) was performed as per the manufacturer’s instructions using RNAScope® Multiplex Fluorescent Reagent Kit v2 for Fixed Frozen Tissue using catalog probes TNF-α (Cat No. 311081), P2Y_12_R (Cat No. 317601-C2) and P2X_7_R (Cat No. 316311-C2).

Briefly, brain sections were heated for 30 minutes at 60°C in a Rototherm Mini Plus (Benchmark Scientific H2024) and post-fixed in prechilled 10% NBF for 15 minutes at 4°C. The tissue was then dehydrated by running it through a 50%, 70% and 100% ethanol gradient for 5 minutes each and then air-dried. Subsequently, after the addition of RNAScope® hydrogen peroxide (incubation for 10 minutes) and washes, target retrieval was performed by boiling the slides for ∼7 minutes in a Bella Food steamer with the target retrieval agent. Afterwards, a hydrophobic barrier was applied (ImmEdge™, Vector Laboratories H-4000), following which sections were incubated in Protease III for 30 minutes at 40°C in a water bath. Probe hybridization (2 hours) was performed, followed by hybridization of AMPs and Opal™ Dyes 520 and 690. Slides were washed twice each, with wash buffer and 1X PBS subsequently, and immunohistochemistry was then performed.

Brain sections were stained with an IBA1 (ionized calcium-binding adaptor molecule 1) primary antibody (microglia/ macrgophage marker, 1:500 dilution, Novus Biologicals NB100-1028) overnight at 4°C, in 0.5% Triton-X (Sigma Aldrich, T8787) on an orbital shaker. The following morning, after two washes with PBS, slides were stained with secondary antibody Alexa Fluor donkey anti-goat 546 for 2 hours, at RT on the orbital shaker. Further, slides were counterstained with DAPI (Advanced Cell Diagnostics), and mounted with Prolong Gold antifade reagent (Molecular Probes) to image on Leica TCS SP8 X White Light Laser Confocal Microscope with a 40x/1.3 oil CS2 objective. Microscope settings including laser intensity, photomultiplier and offset were optimized to yield the highest signal-to-noise ratio, reduced saturated pixels across samples and once finalized, the parameters were held constant between samples. Five regions-of-interests (1 µm z-stack each) in the cortex were imaged in every mouse brain slice, mounted as duplicates. Positive (PPIB) and negative (dapB) control probes were used to confirm the specificity of the probes.

#### Quantification/Data analysis

z-stack images were first pre-processed on the Imaris software (version 10.1.0; Bitplane AG), using a Gaussian filter. Using the “Spots” feature on Imaris, spots were generated for FISH (RNAScope) probes (refer to Supplementary Table 1 for settings). Using the “Cell feature” and customized settings, IBA1^+^ cells were delineated. RNA “Spots” were imported into the “Cell” feature as vesicles, and all quantification of “Spots” and “Cells” were exported using the Imaris “statistics” feature. To have an unbiased approach, the settings for “Spots” for various RNAScope channels were finalized without enabling the IBA1 channel, and vice versa. The “Spots” settings were held consistent between TMEV and PBS brain sections. The parameters for defining “Cells” in the TMEV and PBS conditions varied slightly (Table 1) due to changes in IBA1^+^ expression (Loewen et al., 2016).

### Statistics

Using GraphPad Prism version 10.1.2, based on normality analysis, for comparisons between two groups, the student’s t-test (two-tailed) or non-parametric, Mann-Whitney test was used. Data is represented as mean ± standard deviation (STD) and a p-value of <0.05 was considered statistically significant.

## RESULTS

Mice (10-20 weeks) were intracranially injected with TMEV or PBS and monitored and scored twice a day from days 3-7 post-injection for handling-induced behavioral seizures. Only mice in the TMEV treatment group that had a seizure above grade 3 on the Racine scale were used for subsequent experiments. The timeline for tamoxifen administration, TMEV injection, seizure monitoring and subsequent experiments are shown in Figure 1.

### Reactive microglia respond to exogenous application of damage signals following TMEV infection (5dpi) in the cortex

In TMEV-infected mice, seizures generalize and spread secondarily into the cortex following TMEV infection (Patel et al., 2017). Additionally, microglia in the cortex following infection become reactive (Bell et al., 2020; Loewen et al., 2016). We have demonstrated that reactive microglia in the hippocampus of TMEV-infected mice have a diminished calcium response to exogenous application of ATP and ADP and this is likely due to the decreased expression of P2Y_12_R observed following infection (DePaula-Silva et al., 2019; Wallis et al., 2024). ADP is the primary ligand which binds to P2Y_12_R. ATP signaling has also been shown to lead to cationic influx (including Na^+^, Ca^2+^) through the ionotropic purinergic receptor, P2X_7_R (mechanistic biomarker of epilepsy) and evokes extrusion of K^+^ currents, which in turn has been shown to trigger apoptotic cascades via the NLRP3 inflammasome (Di Virgilio et al., 2017; Muñoz-Planillo et al., 2013). Therefore, we hypothesized that reactive cortical microglia would also have diminished calcium responses following exogenous application of ATP and ADP.

In order to evaluate changes in microglial responses to purinergic damage signals in the cortex as a result of TMEV infection, either ATP (100 uM) or ADP (100 uM) was applied extracellularly to acute brain slices while imaging microglia in mice expressing tdTomato and GCaMP6f. Figure 2A shows panels of the field of view in the respective brain slices (PBS and TMEV) at the instance of the ATP puff. Figure 2B shows time instances corresponding to pre-, during and post-puff time points, and white arrows indicate examples of microglia taken into consideration for calcium fluorescence analysis. The changes in ΔF/F_0_ of GCaMP6f fluorescence are shown in PBS (Figure 2C) and TMEV (Figure 2E) conditions. Upon puffing 100 μM ATP in brain slices from PBS injected and TMEV-infected mice, despite the presence of reactive microglia in the cortex following TMEV infection, their calcium response (ΔF/F_0_) to ATP was unaltered at the peak period of TMEV infection, at 5 dpi (Figure 2 D). Similarly, Figure 2E shows panels of the field of view in the respective brain slices (PBS and TMEV) following the ADP puff. Figure 2F shows panels elucidating pre-, during and post-ADP puff, and panels G and I are representative ΔF/F_0_ traces of microglia as indicated by white arrows in Panel F. The calcium responses of TMEV-infected mice microglia were comparable to those of PBS controls upon 100μM ADP puffing (Figure 2H). These results, along with glial reactivity profiles from previous studies from the lab (Bell et al., 2020; Loewen et al., 2016) suggest that even though there is microglial reactivity in the cortex of TMEV mice, they are able to respond to purinergic damage signals, comparable to PBS controls.

**Figure 2.**
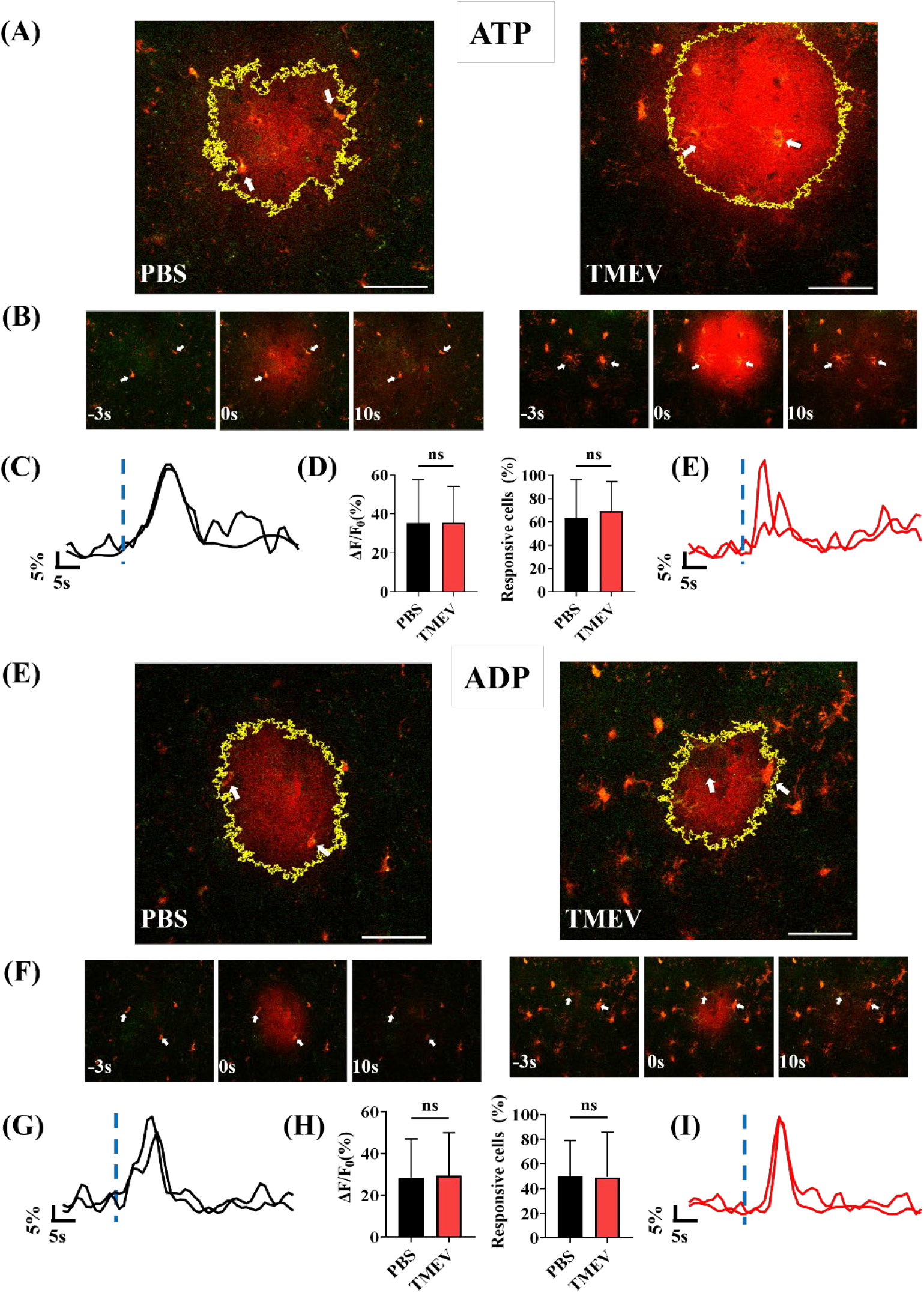
Microglial calcium responses to exogenous application of damage signals (ATP, ADP) remain unchanged as compared to PBS controls, at 5dpi following TMEV infection in the cortex. **(A)** GCaMP6f-expressing microglia respond to an extracellular application of 100 μM ATP in acute brain slices from PBS (left panel) and TMEV mice (right panel). The highlighted yellow area indicates the spread of the ATP puff. **(B)** Images showing the imaging field-of-view at pre (−3s), during (0s) and post puff for PBS and TMEV. White arrows are indicative of example microglia in the given field-of-view. **(C), (E)** Sample ΔF/F_0_ traces of PBS (black) and TMEV (red) of the microglia pointed out in panel B. Blue dashed vertical line indicating time of puff. **(D)** Bar graphs showing maximum ΔF/F_0_ changed and percentage of responsive cells across PBS and TMEV. PBS: n = 6 mice, 12 slices, 95 microglia/ cells. TMEV: n = 4 mice, 7 slices, 56 microglia/ cells. **(E)** GCaMP6f-expressing microglia respond to an extracellular application of 100 μM ADP in acute brain slices from PBS (left panel) and TMEV mice (right panel) with a fluorescent transient. Highlighted yellow area indicates the spread of the ADP puff. **(F)** Images showing the imaging field-of-view at pre (−3s), during (0s) and post puff for PBS and TMEV. White arrows are indicative of example microglia in the given field-of-view. **(G), (I)** Sample ΔF/F_0_ traces of PBS (black) and TMEV (red) of the microglia pointed out in panel F. Blue dashed vertical line indicating time of puff. **(H)** Bar graphs showing maximum ΔF/F_0_ changed and percentage of responsive cells across PBS and TMEV. PBS: n = 6 mice, 11 slices, 72 microglia/ cells. TMEV: n = 4 mice, 8 slices, 40 microglia/ cells. Independent samples t-test or Mann-Whitney test were applied based on normality testing (Shapiro-Wilkins test). Scale bar: 50 µm.

### Differential pro-inflammatory cytokine and chemokine responses in the cortices and hippocampi of TMEV mice

In order to explore brain-region based differences in cytokine and chemokine profiles as a result of TMEV infection, a comprehensive pre-defined inflammation panel (LEGENDplex™ Mouse Inflammation Panel, Biolegend 740446) was conducted to assess protein levels. We found a significant increase in the pro-inflammatory cytokines TNF-α and IL-1α (produced by macrophages and monocytes, which can further activate TNF-α), and the chemokine MCP-1/ CCL2 (regulator of migration and infiltration of monocytes/ macrophages) in the cortex of TMEV-infected mice, as compared to PBS controls (Figure 3A; compare TMEV_Ctx vs. PBS_Ctx). These levels were further increased in the hippocampus (Figure 3A; compare TMEV_HC vs TMEV_Ctx) which represents the primary site of TMEV damage possibly due to TMEV tropism for the pyramidal neurons in this area (K.-A. A. Stewart et al., 2010; K. A. A. Stewart et al., 2010). Additionally, we found a significant increase in levels of the pro-inflammatory/ pro-convulsant cytokine IL-6 in the hippocampus of TMEV-infected mice, as previously described (Figure 3B).

**Figure 3.**
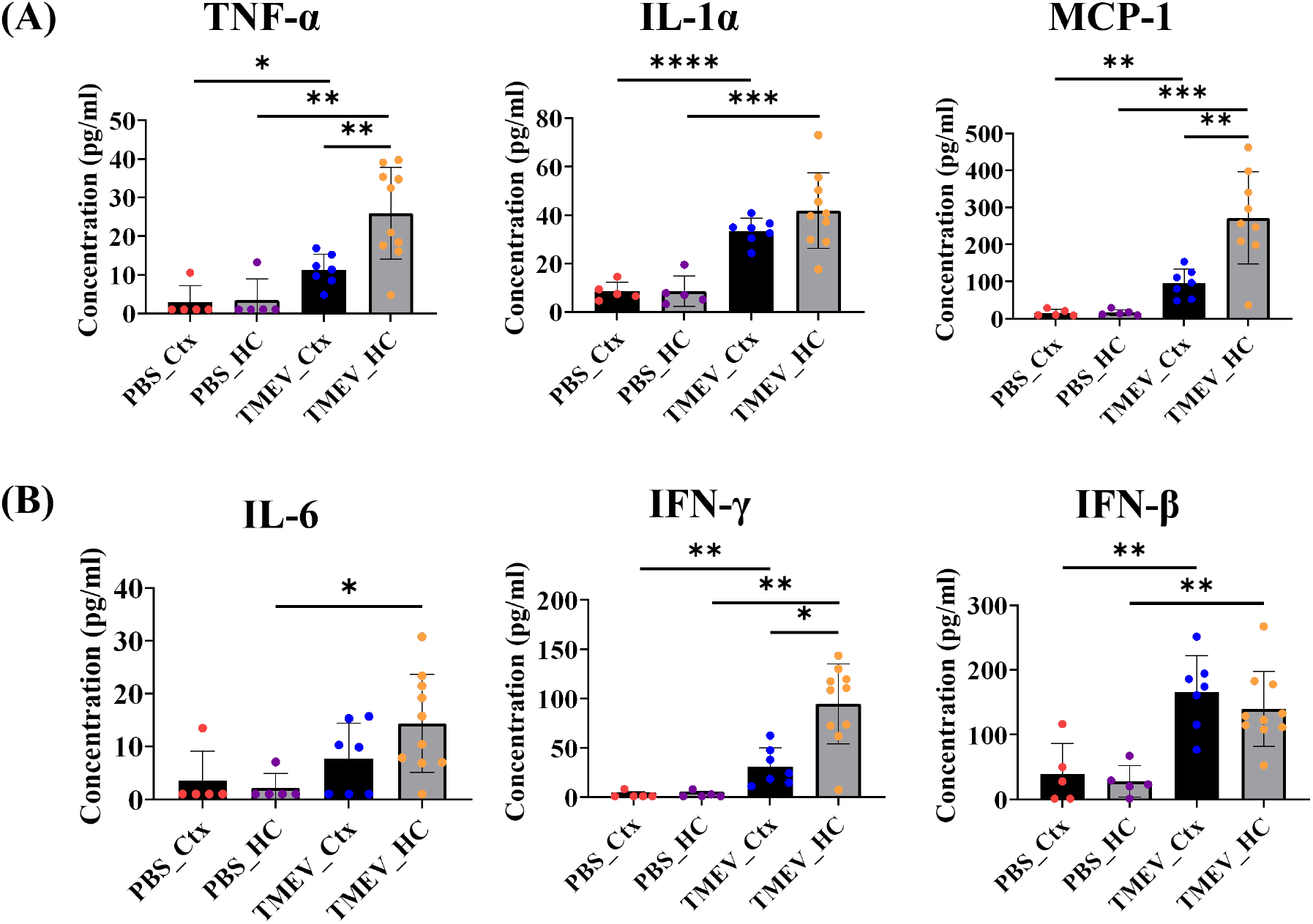
Significant increase in protein levels of inflammatory markers in the cortex and hippocampus of TMEV-infected mice during the acute seizure phase. (A) TNF-α, and the chemokine MCP-1 are increased significantly in TMEV, as compared to PBS controls in both the cortex and hippocampus. IL-1α is increased because of TMEV, and comparable in the TMEV-infected cortex and hippocampus. (B) IL-6 is significantly increased in the hippocampus in TMEV, and IFN-γ and IFN-β are increased in the hippocampus and cortex as a result of TMEV infection, as compared to PBS controls. n = 5 mice (PBS), 10 mice (TMEV). Independent samples t-test or Mann-Whitney test were applied based on normality testing (Shapiro-Wilkins test). ***p<0.001, **p<0.01, *p<0.05.

We also found that the levels of IFN-γ, which is involved in the antiviral response (with ability to inhibit viral replication) and IFN-β, (which reduces excessive neuroinflammation) were increased in both the hippocampus and cortex of TMEV-infected mice compared to PBS controls (Figure 3B). The levels of IL-1α (Figure 3A) and IFN-β (Figure 3B) between the hippocampus and cortex of TMEV-infected mice were comparable for these cytokines. Additionally, we also observed a reduction in levels of IL-23 (inflammatory cytokine for T Helper cell maintenance and expansion), IL-1β (produced by activated macrophages, mediator of inflammatory responses) and IL-27 (may have pro- or anti-inflammatory responses depending on local cues and can lead to IL-10 expression) in the hippocampus in TMEV as compared to PBS controls (Supplementary Figure 1). Overall, for the first time, a gradation of changes in the cytokine and chemokine profiles of cortices and hippocampi were observed in TMEV mice, and in general, there were higher levels of pro-inflammatory cytokines in the hippocampi> cortices> PBS controls. This suggests that there is a heterogeneity of cellular reactivity profiles with respect to pro- and anti-inflammatory cytokines and chemokines as a result of TMEV infection in brain regions crucial for studying viral-infection-induced TLE in mice.

### Increases in purinergic receptor expression and TNF-α following TMEV infection in cortical microglia

In order to study a few downstream receptors of ADP and ATP, we evaluated the purinergic receptor expression of P2Y_12_R and P2X_7_R, along with the inflammatory cytokine TNF-α as a positive control in microglia using RNAScope in situ hybridization and immunohistochemistry to quantify RNA puncta in IBA1^+^ cells. P2X_7_R has been shown to be a biomarker of epilepsy, its hyperactivation has been observed in various disorders, and plays a crucial role in amplifying CNS damage in neurodegenerative diseases (Engel, 2023; Ribeiro et al., 2021). P2Y_12_R plays a critical role in microglial homeostasis, in microglial process extension and retraction (Gómez Morillas et al., 2021). The number of P2X_7_R, TNF-α, and P2Y_12_R RNA puncta were significantly increased in IBA1+ cells in the cortex as a result of TMEV infection at 5 dpi (Figure 4). Collectively, this indicated that there was upregulation of pro-inflammatory markers like TNF-α in the cortex, also observed by our group in the hippocampus (Hanak et al., 2019; Patel et al., 2017; Wallis et al., 2024). This is also consistent with the responsiveness of microglia to purinergic agonists, as observed using calcium imaging in our slice experiments, wherein microglia are as responsive in the cortex following TMEV infection and comparable to that observed in slices obtained from PBS controls. However, P2Y_12_R expression in the cortex is in contrast with what was observed in the hippocampus of TMEV-infected mice during the peak period of infection-where lower expression was concurrent with lower motility to damage signals (Wallis et al., 2024).

**Figure 4.**
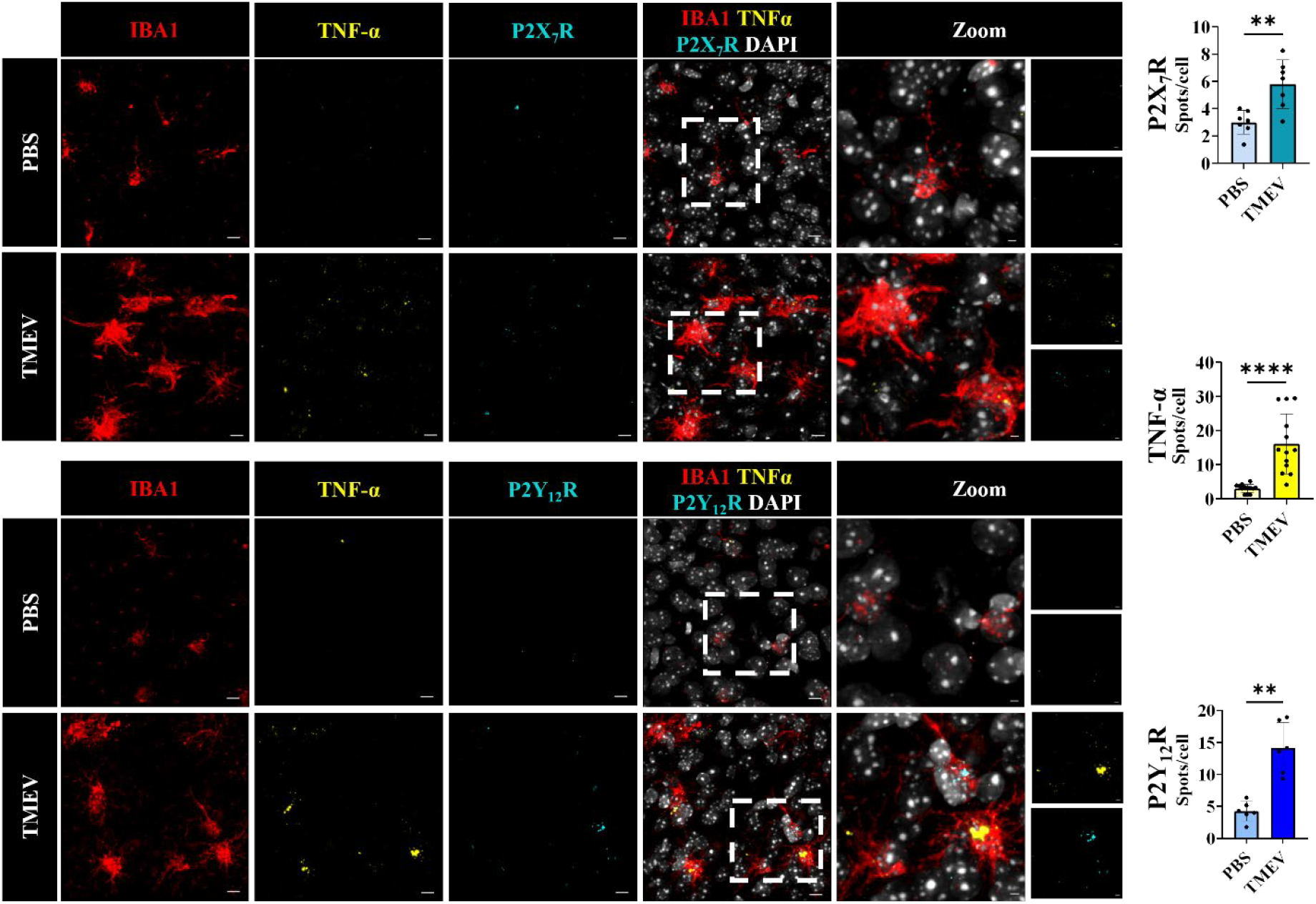
Significant increases in microglial RNA levels of purinergic receptors and TNFα as a result of TMEV infection (acute phase). Increased colocalization of TNFα and P2X_7_R (*in situ* hybridization) (top panel) and TNFα and P2Y_12_R mRNA expression (bottom panel) in IBA1-positive cells (IHC) in the cortex of 5 days post-TMEV infection. Mice: P2X_7_R - n=4 per condition (PBS/ TMEV), sections = 2 per mouse; P2Y_12_R - n=3 per condition (PBS/ TMEV), sections = 2 per mouse; TNFα - n=7 per condition (PBS/ TMEV), sections = 2 per mouse. Independent samples t-test or Mann-Whitney test were applied based on normality testing (Shapiro-Wilkins test). * p<0.05, ** p<0.01, *** p<0.001, **** p<0.0001. Scale bar: 10 µm; for zoomed sections - 3µm.

## DISCUSSION

In this study, we used a multidisciplinary approach to investigate brain region-based changes in microglial reactivity profiles following TMEV infection, at the RNA, cytokine (protein) and cellular calcium imaging scales/ levels. We demonstrated that post-TMEV infection, cortical microglia retain their ability to respond to localized purinergic insults (ADP/ATP), comparable to PBS controls, despite being reactive, in acute brain slice preparations. We also found that the levels of several pro-(TNF-α, IL-1α, IL-6 and IFN-γ) and anti-inflammatory cytokine (IFN-β) are brain-region dependent in TMEV-infected mice, with most increasing in the order of PBS controls, cortices, and hippocampi of TMEV mice. We also observed increases in expression of purinergic receptors crucial for microglial motility (P2Y_12_R) and inflammation (P2X_7_R) in the cortex using RNAScope-FISH. Overall, following TMEV infection, region-based differences demonstrate that reactive cortical microglia retain their ability to respond to purinergic damage signals, the cortex has a higher production of pro-inflammatory cytokines, and there is an increased expression of certain purinergic receptors, as compared to PBS controls.

Microglia are immune sentinels in the brain and form an integral functional unit in the CNS via the ‘quad-partite synapse’ (Schafer et al., 2013) with neurons and astrocytes. Microglia constantly survey the local environment in the brain and play critical roles in responding to CNS insults, brain damage repair, surveillance, neuroinflammation and synaptic pruning (Eyo et al., 2017; Nayak et al., 2012). The dynamic crosstalk between microglia and neurons is facilitated by fractalkine signaling and soluble signaling molecules like ATP act as a “find-me” distress signal for microglial motility and process elongation (Eyo et al., 2014). Particularly, P2Y_12_R is a G_i_ coupled GPCR, which when activated induces chemotaxis in microglia. ADP acts as a primary agonist for P2Y_12_R, while ATP, upon hydrolysis to ADP can serve as an agonist as well. P2Y_12_R plays a crucial initiating role in microglial motility and migration towards CNS injury, surveillance and in extension and retraction of microglial processes (Gómez Morillas et al., 2021; Haynes et al., 2006). Increased expression of purinergic receptors like P2X_7_R and P2Y_12_R have been reported as a result of viral infections in the CNS, and subsequent exacerbation due to production of pro-inflammatory cytokines (Alves et al., 2020). Infected neurons have been shown to recruit P2Y_12_R-positive microglia during encephalitis in the human brain and severity of the infection dictated the level of leukocytes and microglial reactivity/ responses (Fekete et al., 2018). There is reduced microglia-neuron interactions in mice lacking P2Y_12_R, along with increased severity and lethality during status epilepticus (Badimon et al., 2020; Eyo et al., 2014). The implications of increases in P2Y_12_R are still under investigation, while certain groups have reported increases in P2Y_12_R due to viral infections like HIV, as summarized by Alves *et al* (Alves et al., 2020). Studies have also reported decreases in P2Y_12_R due to increased reactivity (Gómez Morillas et al., 2021; Wang et al., 2023). Interestingly, Irino *et al* (Irino et al., 2008) showed that increases in [Ca^2+^]_i_ by P2Y_12_ receptor-mediated PLC activation has been shown to be necessary for ADP-induced chemotaxis of microglia.

In our recent study, decreases in levels of P2Y_12_R and reduced microglial motility to injury (laser burn) were observed in the hippocampus in TMEV-infected mice (Wallis et al., 2024). There is considerable neuronal degeneration of CA1 pyramidal cells within the hippocampus of TMEV-infected mice and a downregulation of P2Y_12_R was also reported in bulk RNA-seq experiments by DePaula-Silva *et al* (DePaula-Silva et al., 2019) as a result of TMEV. Interestingly, in the present study, we show that reactive cortical microglia in TMEV-infected mice have increased P2Y_12_R expression at the RNA level (Figure 4) at the same time point post infection that microglia in the hippocampus exhibit a decreased expression (Wallis et al., 2024). The present experiments evaluating cytokine panels (Figure 3) also corroborates the increases in expression of several pro-inflammatory cytokines including TNF-α, IFN-γ and IL-1α, in the cortex of TMEV-infected mice versus PBS controls. Interestingly, the levels of some of these cytokines and the chemokine MCP-1 are further increased in the hippocampus, in accordance with previous studies from the lab detailing increased pro-inflammatory profiles in the hippocampus of TMEV mice (Bell et al., 2020; Loewen et al., 2016; Patel et al., 2017). It will be important to assess microglial motility in the cortex following TMEV infection to determine if microglia in the cortex retain their motility. Given that there is an increase in the mRNA expression of P2Y_12_R in the cortex and robust purinergic responses to agonist applications, we would hypothesize that motility is also unaffected. The therapeutic value of P2Y_12_R (Chen et al., 2019) also implores further assessment of implications of its beneficial effect in the cortex, as observed in our results in TMEV-infected mice.

Our data shows an upregulation of P2X_7_R at the RNA level, at the peak period of TMEV infection (5 dpi) in the cortex, as compared to age-matched PBS controls. This further defines the reactivity of IBA1+ cells in the cortex of TMEV mice, but further evaluation at various extended time points (like 14 dpi) could shed light on the evolution/ reduction of these reactivity profiles. The P2X_4_ receptor is also stimulated by ATP in microglia and is involved in chemotaxis along with P2Y12_R_ (Ohsawa et al., 2007). Another crucial purinergic receptor to be investigated in TMEV is the P2Y_6_ receptor. UDP activation of P2Y_6_R leads to microglial phagocytosis of dying cells in epilepsy, and future studies could help unravel the phagocytic changes in purinergic signaling as a result of TMEV infection across the cortex and hippocampus.

P2X_7_R, an ionotropic purinergic receptor has been shown to be a mechanistic biomarker in epilepsy, and other conditions like Alzheimer’s Disease and COVID-19 (Engel, 2023; Illes, 2020; Ribeiro et al., 2021). As a result of ATP signaling, the P2X_7_ receptor allows for an influx of cations, including Ca^2+^ and Na^+^, and evokes extrusion of K^+^ currents. P2X_7_R signaling is also crucial to the NLRP3 inflammasome-mediated triggering of apoptotic cascades, leading to cell inflammation and cell death (Di Virgilio et al., 2017). Furthermore, microglial plasma membrane blebbing can be a consequence of ATP-induced P2X_7_R activation (Illes, 2020). Potassium efflux has been shown to trigger activation of caspases and inflammasome activation via the P2X_7_ receptor (Muñoz-Planillo et al., 2013). Recently, it has been discovered that K^+^ efflux also mediates P2Y_12_R-dependant inflammasome activation (Suzuki et al., 2020), thereby unraveling novel roles of these receptors.

In conclusion, we evaluated region-specific differences in cytokine profiles and determined the microglial purinergic receptor-mediated calcium imaging responses in the cortex in the TMEV model of infection induced TLE. Despite microglia being reactive in the cortex (confirmed by P2X_7_R increases, phenotypic morphology, and cortical increases in cytokine expression), there are no changes in the calcium signaling response to application of ATP or ADP following infection. Future work investigating 1) different time points, like 2 dpi (before seizures begin in the TMEV model) and 14 dpi (after seizures resolve and the virus is cleared in the TMEV model), 2) microglial motility in the cortex of TMEV-infected mice and 3) an in-depth study of other purinergic receptors using tools like spatial transcriptomics, could unravel more information about the evolution of the neuroinflammatory states in cortical microglia as a result of TMEV infection.

## ABBREVIATIONS

2-P: Two-photon
3D: Three dimensional
°C: Degree Celsius
A3: A3 adenosine receptor
A2A: A2A adenosine receptor
aCSF: artificial cerebrospinal fluid
ADP: Adenosine diphosphate
AMP: Adenosine monophosphate
ATP: Adenosine triphosphate
[Ca^2+^]: Calcium
CNS: central nervous system
CreERT2: Cre recombinase – estrogen receptor T2
CX3CR1: C-X3-C Motif Chemokine Receptor 1
ΔF/F_0_: Change in fluorescence intensity relative to the baseline fluorescence intensity
dpi: Days post-infection
FISH: fluorescent *in situ* mRNA hybridization
GPCR: G-Protein coupled receptor
Hz: Hertz
IACUC: Institutional Animal Care and Use Committee
IBA1: Ionized calcium-binding adapter molecule 1
IFN-β: Interferon-Beta cytokine
IFN-γ: Interferon-Gamma cytokine
IL-1α: Interleukin 1-alpha cytokine
IL-6: Interleukin-6 cytokine
IL-10: Interleukin-10 cytokine
IL-23: Interleukin-23 cytokine
IL-27: Interleukin-27 cytokine
i.p.: Intraperitoneal injection
JAX lab: Jackson Laboratory
Lck-GCaMP6f: genetically encoded fast calcium indicator tethered to the membrane
MCP-1: Monocyte chemoattractant protein-1
mOsm: milliosmole
Min: Minute(s)
mL: Milliliter
mm: Millimeter
mM: Millimolar
mRNA: Messenger ribonucleic acid
ms: Millisecond
mW: Milliwatt
MΩ: Megaohm
NA: Numerical aperture
NBF: Neutral buffered formalin
NLRP3: Nucleotide-binding domain, leucine-rich–containing family, pyrin domain–containing-3
nm: Nanometer
PCR: Polymerase Chain Reaction
P2XR: P2X purinergic receptors
P2X_7_R: Purinergic receptor P2X_7_
P2Y_12_R: Purinergic receptor P2Y12
P2RY: P2Y purinergic receptors
PBS: Phosphate-buffered saline
PFU: Plaque-forming units
PLC: Phospholipase-C
PSI: Pounds per square inch
RNA: Ribonucleic acid
RO: Reverse Osmosis
ROI: Region of interest
s: Second(s)
RT: Room temperature
STD: Standard deviation
SA-PE: Streptavidin-phycoerythrin
TAM: Tamoxifen
tdTomato/ tdTom: tdTomato fluorescent protein
TLE: Temporal lobe epilepsy
TMEV: Theiler’s murine encephalomyelitis virus
TNF-α: Tumor necrosis factor alpha
µg: Microgram
µL: Microliter
µm: Micron/micrometer
µM: Micromolar

## ACKNOWLEDGEMENTS

This work was supported by NIH/ NINDS, R37NS065434 (KSW) and NIH/D-SPAN F99NS125773 (LAB). The authors would like to thank E. Jill Dahle, M.S. Alexandra Petrucci, Ph.D., and Carolina Moncion, Ph.D. for their technical assistance and insightful discussions. We also thank the Health Science Center Cell Imaging Core facility and its staff, Xiang Wang, Ph.D., and Anton Lassen, Ph.D., for their assistance with software, microscopy and IMARIS platforms.

