## Supplementary Table 1 and Supplementary Figure 1 for "Investigating the role of cortical microglia in a mouse model of viral infection-induced seizures"

**Supplementary Table 1: Parameters used for RNAScope analysis**

| **Spot detection** | **Values** |
| --- | --- |
| Segment only Region of Interest | No |
| Different spot sizes (region growing) | No |
| Classify spots | No |
| Object-Object statistics | Yes |
| Estimated XY Diameter | 0.8 um |
| Enable shortest distance | Yes |
| Model PSF-elongation along z axis | No |
| Background subtraction | True |
| Filter (Quality) | > 650 (TNFα),  > 800 (P2X_7_R and P2Y_12_R) |

| **Cell detection** | **PBS** | **TMEV** |
| --- | --- | --- |
| Segment only ROI | No | |
| Select Cell detection type | Cell body labeling | |
| Smooth | Yes | |
| Filter Width | 0.426 um | |
| Background Subtraction | No (Done during pre- processing) | |
| Cell Manual Threshold | 8000 | 9000 |
| Split Touching cells  (by seed points) | Yes | |
| Nuclei options | No | |
| Diameter of Seed points | 15 um | |
| Filter Cells (Cell volume) | > 300um^3^ | > 600um^3^ |

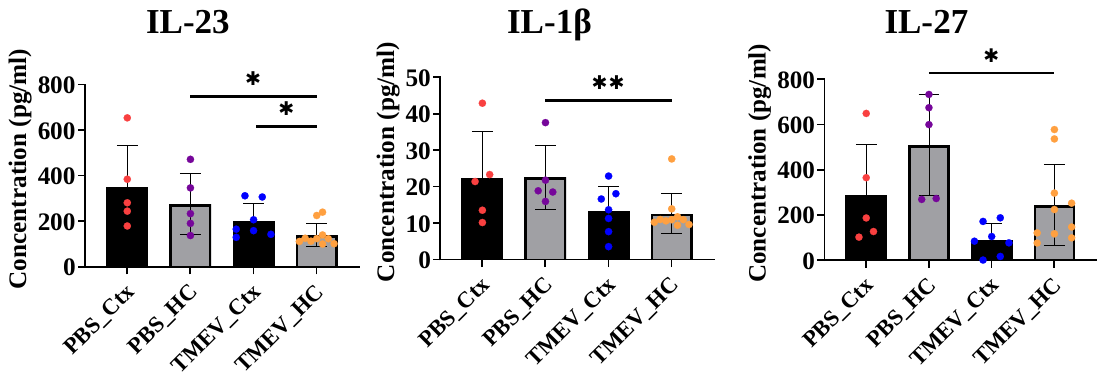
**Supplementary Figure 1. Significant decreases in certain cytokine levels in the hippocampus, due to TMEV infection, during the acute seizure phase.** Protein levels of IL-23, IL-1β and IL-27 were significantly decreased in the hippocampus of TMEV-infected mice, as compared to PBS controls. n = 5 mice (PBS), 10 mice (TMEV). Independent samples t-test or Mann-Whitney test were applied based on normality testing (Shapiro-Wilkins test). **p<0.01, *p<0.05.
